# Automated Facial Landmark Analysis vs. Manual Coding: Accuracy in Dog Emotional Expression Classification

**DOI:** 10.1101/2025.10.23.683931

**Authors:** Jiao-Ling Appels, George Martvel, Anna Zamansky, Stefanie Riemer

## Abstract

Emotional expression in dogs is central to dog-human interactions. Reliable indicators are essential for interpreting animal emotions; however, their predictive value remains debated. Fireworks, which elicit fear in many dogs, provide a real-world context for examining affective states. We used machine learning to classify firework vs. non-firework situations from dogs’ behaviour and expressions using two approaches: (1) ethogram-based manual coding of behaviours and (2) automated analysis of facial landmarks. The Random Forest model based on manual coding achieved a high accuracy (0.83) and 1.0 predictive validity, identifying backwards-directed ears and blinking as key fear indicators. The best automated facial landmark model reached up to 0.80 accuracy and 0.77 predictive validity. Manual coding performed better, likely due to richer semantic content and full-body observations. Our findings demonstrate that machine learning can classify canine emotional states from both manually coded and automated methods, offering potential for future automated welfare monitoring.

**Highlights:**

- Machine learning approaches classify firework-related fear in dogs reliably
- Ear position and blinking are key indicators
- Ethogram-based coding yielded stronger predictions than automatically detected facial landmarks
- Automated landmark analysis shows potential for scalable welfare monitoring

Graphical abstract

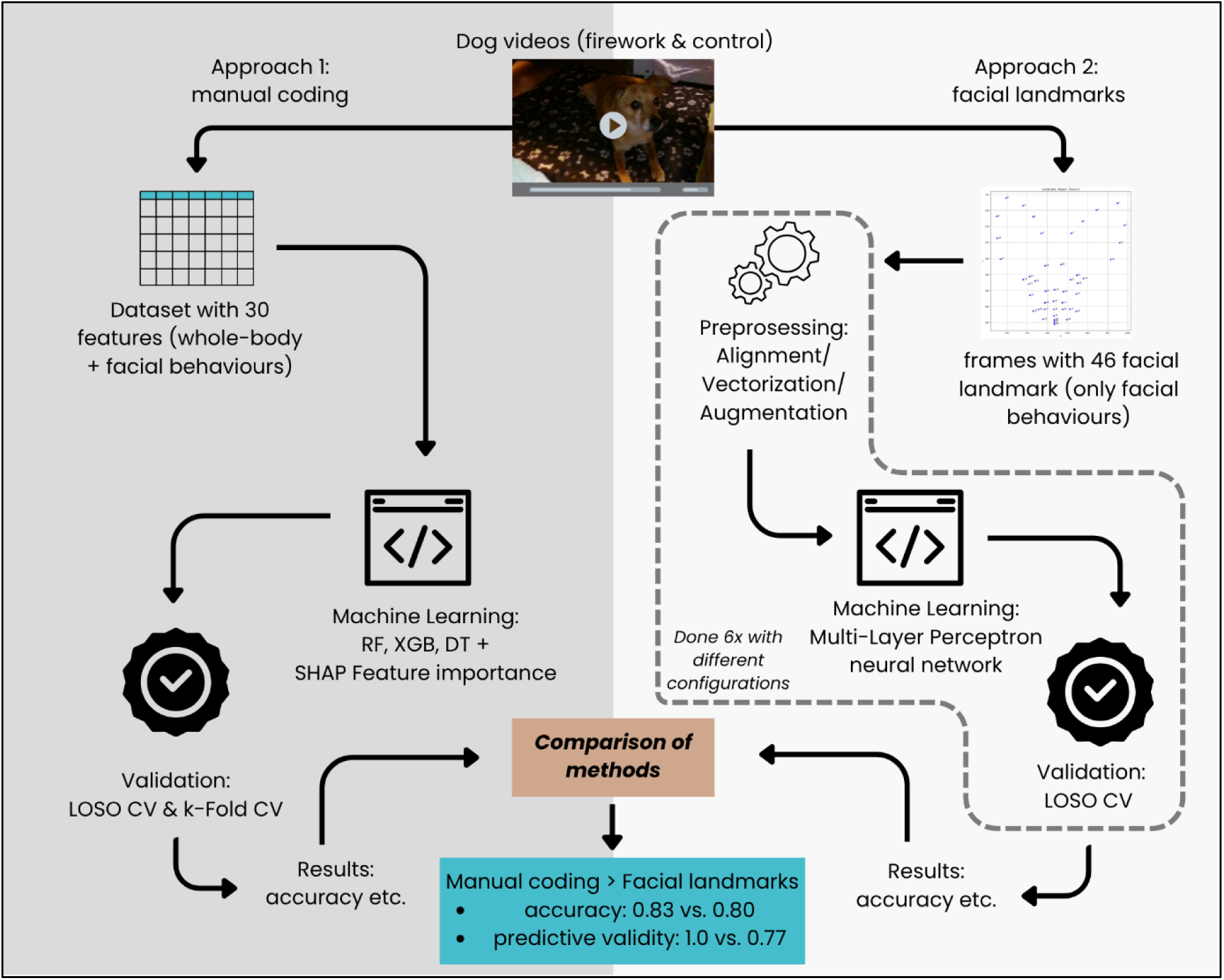

**Subject areas:** Emotional expression · Behavioural classification · Machine learning · Facial landmarks · Facial expressions

## Introduction

There is increasing interest in objectively assessing companion animal welfare^1,2^, aiming for both minimising negative states such as fear and increasing positive emotions such as happiness^3^. Accurate indicators of emotional states in nonhuman animals are essential; however, drawing inferences from behavioural and/or physiological indicators remains challenging due to variations in context, action tendencies, and behavioural expressions^4^. Many dogs are highly sensitive to noises such as gunshots, thunderstorms, and fireworks^5,6^, with nearly half showing fearful reactions specifically to fireworks^7–10^. Common fear-related behaviours include hiding, panting, trembling, and barking^7^, while more subtle cues—such as freezing, blinking, and ear position^8^—may often go unnoticed. While fear responses are adaptive behaviours that help animals avoid potential threats, in dogs with heightened sensitivity, these responses can be severe and prolonged, significantly impacting their welfare^11,12^. To enhance welfare assessment, valid, reliable, and robust emotional indicators are needed^13^.

To make inferences about likely affective states animals may experience, we can draw from (1) cognition: changes in attentional, perceptual, and inferential processes (appraisals), which can often be inferred by animals’ choices and behaviour^3,14^, (2) physiology: neural activation and associated neuroendocrine responses^3,14^, (3) behaviour/action tendency: predisposition for performing specific responses^3^, and (4) expression: subtle expressions such as postures, facial expressions, vocalisations and release of odours or pheromones^3,14^. However, some common physiological indicators, such as increases in cortisol, core body temperature or heart rate^15^, may not only reflect arousal associated with negative emotions, but they also occur in positive contexts^16^, such as in relation to food^17^ or sexual interactions^16^. Moreover, a given behaviour does not necessarily have a one-to-one association with a specific emotion^18^. Alternative behavioural strategies may be observed—for example, in a fearful situation, the subject can react with active avoidance (flight or hide), passive avoidance (freeze) or active defence (threat, attack)^18^.

Various approaches can be employed to measure the fourth component, expression, ranging from behavioural ethograms to fine-grained analyses of subtle facial movements to AI analysis of facial or body postures or movements. Ethogram-based coding remains a standard approach in controlled studies. It provides detailed, semantically rich data, but it is labour-intensive and potentially biased^19–21^. Using this method, researchers have observed differences in expression in dogs between contexts, such as between a firework and a non-firework situation^8^, between frustration and positive anticipation^22,23^, and in response to other noises like thunderstorms and gunshots^7^. However, while ethograms capture whole-body behaviour, they offer limited detail on facial expression. The Facial Action Coding System (FACS) represents the gold standard for objectively coding facial expressions based on underlying muscular movements ^24,25^. Originally developed for humans ^24,25^, FACS has been adapted to various animal species to capture species-specific facial expressions, including dogs (DogFACS)^26^. Using DogFACS, it has been demonstrated that different affective states are reliably associated with certain facial expressions in dogs^13,25,27^. However, the disadvantage of this method is that coding is extremely time-consuming and requires high-quality videos. Alternatively, AI approaches are increasingly being used in the analysis of animal (dog) facial emotional expression^24^.

Automated facial landmark analysis is an emerging method for detecting facial expressions. This technique identifies key points on the dog’s face—such as around the eyes, ears and mouth—to capture important morphological details. Changes in the configurations of such landmarks allow possible inferences about emotional states^28,29^. Facial landmark analysis is a non-invasive and objective approach that is especially appealing for automated emotion recognition and welfare monitoring in animals^19,30,31^. In dogs, landmark-based methods have been successfully used to identify behaviours associated with different emotions^22,32,33^. However, dogs can be particularly challenging for automated approaches due to their extreme variability in morphology. For example, when detecting and classifying dogs, pointy-eared dogs are more easily recognised by the model compared to floppy-eared dogs^34^. Despite some issues with using landmark-based approach for emotion categorisation from video recordings, including obscured faces or extreme angles or head rotations, we recently demonstrated for the first time that this method can be successfully applied to an inhomogenous data set collected “in the wild” (owner-recorded videos of dogs during New Year’s Eve fireworks and on a control evening without fireworks): Machine learning successfully differentiated between situations based on automatically detected landmark configurations in the faces of dogs of various breeds and morphologies ^35^. In agreement with the manually coded data^8^, the positioning of the ears differentiated most strongly between the conditions, even though only landmarks from the base of the ears, but not of the ear pinnae, were included, given much variation in ear shape between individuals^35^.

The great advantage of machine learning is that neural networks efficiently analyse large datasets, detecting complex patterns often missed in conventional analysis. Studies focusing on individual behavioural indicators, such as those obtained by ethogram-based or FACS methods, are typically limited in their predictive validity^13,36,37^, as even a statistically significant difference in behaviour does not always mean that the conditions can be accurately predicted from the data^13^. Automated classification systems using machine learning can offer an objective alternative to traditional statistical analysis by utilising manually coded behavioural data^38,39^. Machine learning is ideally suited for analysing complex animal behaviour^25,29,40^. It provides a powerful tool for detecting both overt and subtle fear responses in dogs by analysing multiple behavioural indicators simultaneously^24,32^.

Multiple studies suggest including both statistics and machine learning to establish the accuracy of the method and to make predictions^13,41^. Assessing predictive validity—how well a model predicts future behaviour or outcomes^37^—is essential to ensure that behavioural indicators and model outputs accurately reflect expressions associated with differing emotions, rather than coincidental or superficial patterns^42^. Predictive validity can also be used to compare machine learning methods. To our knowledge, no study to date has compared the accuracy of human-derived ethogram-based coding vs automated emotion detection in identifying different emotional states in dogs, based on machine learning applied to both datasets.

This exploratory study aims to identify candidate behavioural indicators associated with firework-related fear in dogs. Specifically, it evaluates whether machine learning models can classify firework versus non-firework situations based on two distinct data sources: (1) manually coded behavioural features from an ethogram-based video analysis and (2) facial landmark data, obtained via automated AI video analysis. The study further investigates the predictive validity of the most informative behavioural indicators identified by the models. It seeks to explore whether machine learning can accurately classify firework-related fear in dogs using manual behavioural coding or facial landmark data, and which features provide the strongest predictive value. Additionally, we investigate whether key behaviours and facial features retain their predictive power across individuals. This approach may support the development of automated tools for recognising fear and stress in dogs, ultimately enhancing welfare monitoring in real-world settings.

## Results

### Methods summary

We applied two complementary machine learning approaches to classify dogs’ emotional responses to fireworks. The data consist of previously collected videos of dogs filmed during New Year’s Eve and an evening without fireworks. In the first approach, an ethogram-based method was used: coders manually annotated body positions and facial behaviours of dogs (n = 36) according to an established ethogram^8^, which were then analysed using tree-based machine learning models (Random Forest, XGBoost, and Decision Tree). In the second approach, facial expressions were quantified automatically using facial landmark detection, extracting 46 facial points per frame. These frames were analysed using a Multi-Layer Perception neural network under four different preprocessing configurations. This approach also compared between an unbalanced (n = 25 dogs; 32 videos) and a balanced (n = 7 dogs; 14 videos) dataset. Both approaches were evaluated using accuracy, precision, recall, F1-score, and cross-validated using Leave-One-Subject-Out. Together, these methods allowed comparison between a manually coded and an automated system.

### The Random Forest model yields the highest accuracy, with ear position as the key indicator

The Random Forest (RF) model of the manually coded data achieved the highest accuracy (0.80), with *Ear 1* (low, i.e most backwards-directed ears on a 5-point scale) and *blink/min* (frequency of blinking per minute) as the top predictive features. In contrast, the XGB and DT models showed lower accuracies (0.60 and 0.67, respectively) and identified different key features (Figure 1, Table S1). Both RF and DT models ranked Ear 1 (low) as the most important feature.

**Figure 1:**
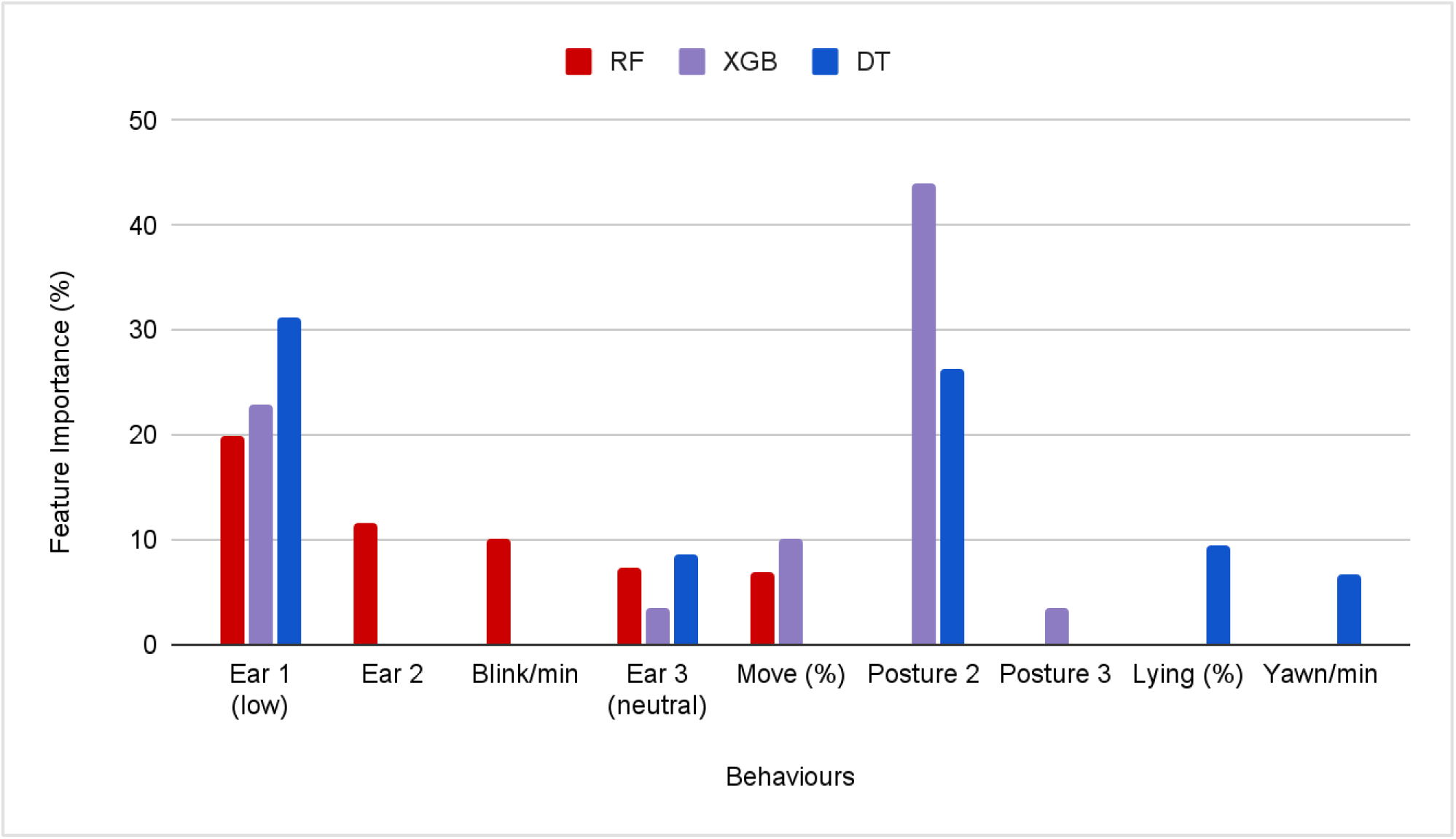
The model performance of the three models and their five most important features.

### Backwards-directed ears and blinking are the best indicators of firework fear

The SHAP analysis reveals each feature’s contribution to predicting firework situations in the Random Forest model (Figure 2). High values of Ear 1 and a high frequency of blinks per minute increase the likelihood that the model identifies a firework event, whereas high values of sleep decrease that likelihood. Comparing RF’s top features with Cohen’s *d* effect size from the manually coded data (Gähwiler et al.^8^; Table S2) confirms ear position as the most distinguishing behaviour. However, RF identified blinking as a key predictor, which was not significant in manual statistics when applying sequential Bonferroni correction^8^.

**Figure 2:**
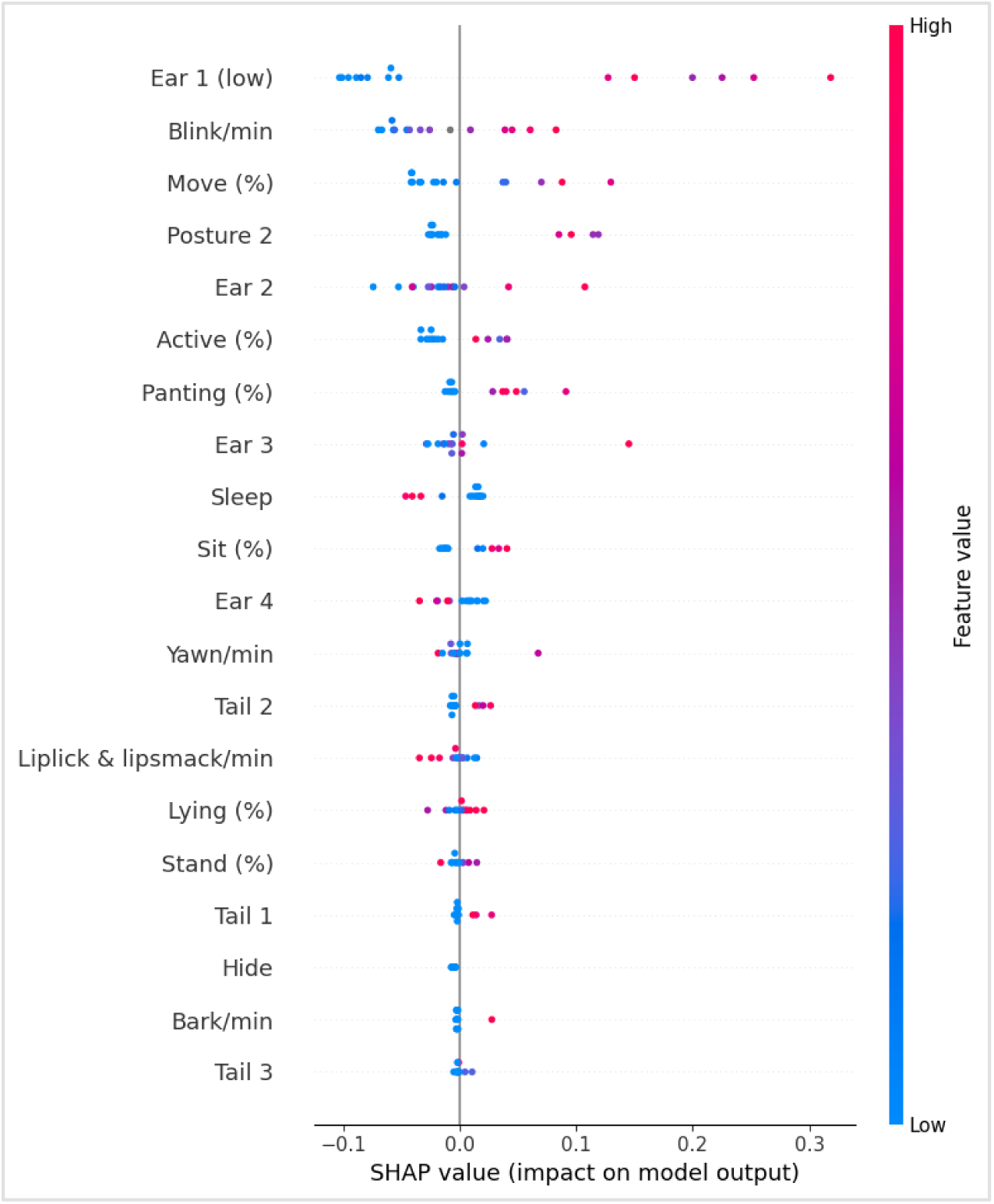
The impact of each feature on the Random Forest model on predicting the firework situation (Table S2).

### Model-selected reliable indicators have an excellent predictive validity

Multiple feature selection strategies were tested and cross-validated before choosing the most predictive set for generalizability. Based on SHAP best features and validation, the selected SHAP features are: all five ear positions, Blink/min, Move (%) & Panting (%). Subject-aware validation methods, “Leave-One-Subject-Out” and “k-folds∼number of subjects/10”, revealed that the model using SHAP or all behavioural features with a LOSO split achieved the highest overall F1 score (0.889) (Table 1). However, the model using a 4-fold subject-based split also yields high accuracy (0.833) and recall (1.0).

**Table 1:** Results of four different predictive validity methods using a different number of selected features^a^.

| Metric | LOSO SHAP | LOSO All | K=4 SHAP | K=4 All |
| --- | --- | --- | --- | --- |
| Accuracy | **0.833** | **0.833** | 0.833 | 0.833 |
| Precision | 0.833 | 0.833 | 0.812 | 0.750 |
| Recall | 1.0 | 1.0 | 0.917 | 1.0 |
| F1 Score | 0.889 | 0.889 | 0.843 | 0.857 |
| ROC AUC | **1.0** | **1.0** | 0.875 | 0.806 |

<sup>a</sup>Best values are in bold.

### Training the algorithm on a balanced dataset yields better classification accuracy despite a lower sample size

A total of 55 videos were excluded from the initial dataset because fewer than 50% of their frames contained detectable landmarks. For the remaining videos, frames were considered “good” if the model’s detection confidence was greater than zero (conf ≠ 0.0), as a score of 0.0 indicates no visible landmarks. The final dataset included 32 dog videos of 25 dogs, comprising 15 videos from non-firework situations and 17 videos from firework situations. Of the 25 dogs, only 7 had videos available for both conditions, allowing for a separate balanced subset analysis. Each subset was analysed under different configurations: 3, 5 or 10 automatically extracted frames per video.

As described in the Methods, each version of the MLP model was trained under different preprocessing conditions (with or without facial landmark alignment and with or without data augmentation). The classification accuracy varied notably depending on the preprocessing pipeline and dataset configuration (Table 2; Table S3).

**Table 2:**
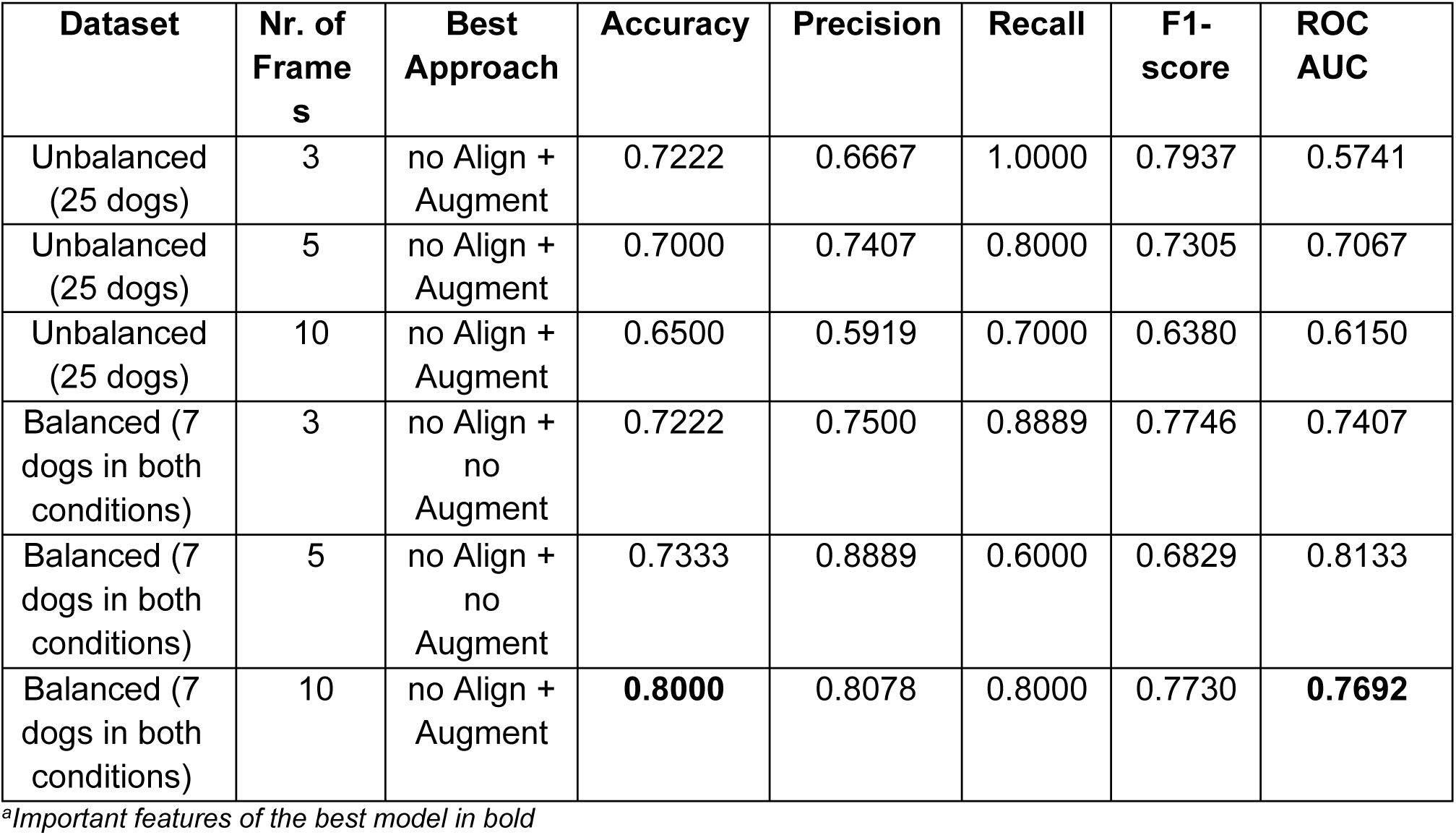
Ablation table of LOSO evaluation results for all configurations^a^.

A comparison revealed that the best results were obtained when using 7 dogs with videos in both conditions and 10 frames per video, without alignment but with augmentation. This setup achieved an accuracy of 0.714 and ROC AUC of 0.816, indicating good generalisation in distinguishing between firework and non-firework expressions. In contrast, models trained on a larger dataset with 25 dogs—but unbalanced in condition (i.e. some dogs featured only in the firework dataset and others only in the control dataset)—performed notably worse across all configurations. Accuracies were typically around 0.55-0.65, with some models performing as low as 0.33, suggesting that imbalanced data limits model performance.

## Discussion

The dogs showed distinct facial and bodily expressions in the two conditions (firework/control), demonstrating that affective states can be reliably deduced from behavioural coding. Indeed, an extremely high predictive power of up to 100% was achieved based on the machine learning analysis of manual coding. Thus, while individual indicators may not allow accurate deductions of affective states on their own^25^, the machine learning approach allows incorporating multiple indicators simultaneously, achieving high predictive validity. Meanwhile, the optimal achieved results were considerably lower when focusing on automatically detected facial landmarks with an accuracy of 0.80 and a predictive validity of 77%.

This is comparable to the results of Boneh-Shitrit et al.^24^, who used supervised machine learning to classify dog faces from a positive and a negative (frustration) condition. Our best tree-based model (Random Forest) reached a higher accuracy (0.80) compared to theirs (0.72). However, their other tree-based models (XGBoost & Decision Tree) performed better (0.71 vs. 0.60/0.67 in the current study). Boneh-Shitrit et al.^24^ also reported slightly higher accuracies (up to 0.82) and higher F1-scores (up to 0.83) using supervised deep learning approaches. This may be explained by their more balanced dataset with fewer subjects that were morphologically similar (Labrador retrievers). Their data was collected under standardised laboratory conditions, and all dogs were videorecorded from the same (frontal) angle. Conversely, the dataset from the current study was very messy, consisting of owner-provided home videos of their dogs of widely varying quality, angles, and visibility of the dogs, and including dogs of various morphologies. Therefore, yielding an accuracy of 0.80 under these conditions is still a remarkable result.

The results of the classification of the manually coded data show that the Random Forest (RF) model outperformed XGBoost (XGB) and Decision Tree (DT), in line with the results of Boneh-Shitrit et al.^24^ This is likely due to RF’s ability to handle non-linearity, reduce overfitting, and capture subtle feature interactions^43,44^. XGB may require more fine-tuning for optimal performance, particularly in tuning hyperparameters^45^, while DT models tend to be less robust due to their tendency to overfit^46^.

Differences in feature prioritisation stem from the models’ evaluation methods: RF and DT use Gini impurity, whereas XGB relies on gain-based evaluations^47^, highlighting how model choice influences feature importance. Unlike the previous study by Gähwiler et al.^8^, which averaged ear, tail, and posture positions, this study considered the five positions as separate features to capture subtle behavioural variations. This approach is in line with Bremhorst et al.^25^, who highlighted that subtle differences in ear movements—such as distinguishing between “Ears rotator” and “Ears flattener”—may reflect distinct emotional states and thus warrant more detailed classification.

Ear positions have been found to be particularly important for conveying emotional states in various species with mobile ears, such as sheep^48^, mice^49^, and cats^30^. Previous studies in dogs have identified distinct ear movements that may differentiate between positive and negative emotional states: the “ears flattener”, “ears rotator”, “ears adductor”, and “ears downwards” actions, as defined in the DogFACS system^13,50^. Ears held upright may signal positive valence (e.g., happiness), but they can also indicate vigilance or heightened arousal in response to potentially fearful or unpredictable situations^8,13,51^. Reefmann et al.^48^ suggested that upright ears in sheep may coincide with negative emotional states, especially when animals anticipate aversive events. In the context of firework-related fear, backwards-directed ear movements—particularly the ear rotator—have been most consistently associated with non-social fear in dogs with erect ears^8^. However, this ear posture is not exclusive to fear and can also occur during frustration or affiliative greeting situations involving active submission^24,25,52^. Despite this variability, ears flattened or turned backwards have been repeatedly reported to be associated with negative emotional states—particularly fear, anxiety, or submission^4,8,33^. Further work on the possible distinct functions of the different ear actions is needed (e.g., Gähwiler et al.^8^).

In contrast to a previous study by Martvel & Riemer^35^, which excluded the ear pinnae landmarks due to the high variability in ear morphology among dogs, this study retained them. The decision to include ear pinnae landmarks was made to enable comparison with the manual behaviour coding approach, where ear position was an important feature. Despite the diversity in breeds and ear types, the classification performance remained high. However, accuracy might have been higher if the dataset had been limited to dogs with erect ears or short hair, given previous findings showing that landmark detection performs better in dogs with upright ears or smooth facial fur, as opposed to those with long hair covering facial features^34^. It also remains possible that excluding the ear pinnae altogether could have improved accuracy by reducing variability of the ear landmarks, considering that the landmarks related to the ear base differentiated most strongly between the fireworks and control conditions in Martvel & Riemer^35^.

The SHAP analysis revealed a key discrepancy with traditional statistical methods. While analysis of manually coded data initially showed significant differences in blinking between firework and non-firework situations, this significance disappeared after sequential Bonferroni correction, which can be overly conservative^53^. However, the fact that blinking was identified as an important feature by the Random Forest model suggests that the result is likely not due to chance^54^. Traditional statistical tests evaluate variables independently, whereas SHAP considers feature interactions and nonlinear relationships, identifying blinking as a key predictor despite its lack of standalone statistical significance^55,56^. This suggests that blinking may meaningfully contribute to the prediction when combined with other behaviours, a factor that may be overlooked in conventional approaches.

The predictive validity of the manual behavioural coding showed that the model using LOSO validation and only SHAP or all behavioural features achieved the highest F1-score. This suggests an optimal performance in balancing false positives and false negatives^57^. This more conservative LOSO method also yielded the highest ROC AUC (1.000), reflecting strong discriminative power and excellent generalizability to unseen individuals^42,46^. Even though the k-fold subject-based split achieved equally high accuracy, the predictive performance is lower. These results illustrate a trade-off between maximising predictive performance and maintaining model interpretability^46,47,58^. While including all features may offer minor improvements in ambiguous cases, the top SHAP-identified indicators (ear positions, blinking, movement, and panting) were sufficient to achieve competitive accuracy and recall across all validation strategies, supporting their use in future predictive tools and welfare assessment frameworks. This result was also found in Boneh-Shitrit et al.^24^, where models considering all variables yield higher evaluation metrics compared to the models with a subset of variables.

Among the landmark-based models, the best performance was achieved using no alignment of facial landmarks but only augmentation, particularly when using the balanced subset (7 dogs; both conditions; 10 frames per video), yielding an accuracy of 0.80 and an F1-score of 0.77. Although alignment preprocessing is assumed to help standardise facial structure by reducing variability^29^, alignment did not consistently improve results in the current study. One possible explanation is that our alignment procedure (eye-based) may have suppressed key emotional cues by removing informative spatial variation, especially around the ear and head position. For instance, aligning based on the eyes may inadvertently shift or distort ear landmarks, which are key to fear expression (e.g., ears rotated backwards).

In contrast, models with alignment and no augmentation generally showed lower performance, suggesting that alignment removed some useful variation related to ear orientation and head pose. Without augmentation, the model also had less diversity to learn from. Similarly, models with alignment and Gaussian noise injection (augmentation) to simulate natural variability in facial expression, using a small standard deviation to avoid large distortions, performed worse. This supports the idea that emotion-related facial expressions in dogs rely on subtle spatial patterns of facial landmarks, which may be disrupted even by minor noise. While noise-based augmentation is commonly used to improve model generalization^28,33^, its effectiveness in this context seems limited, likely due the high sensitivity of the task to precise spatial configurations and possibly the small dataset size.

The current noise-based augmentation method was adapted from Feighelstein et al.^29^ on facial expressions of pain in cats. Our results for fear detection in dogs show comparable performance to this work^29^. While Feighelstein et al.’s^29^ results achieved higher accuracy in all four approaches, we achieved an accuracy of 0.80 in one approach using the same model architecture and LOSO evaluation, despite a smaller dataset. Fear detection in dogs may be inherently more challenging than pain detection in cats, firstly due to greater morphological variations in dog faces^34^, and because fear expressions may be variable across individuals (c.f. Gähwiler et al.^8^). Additionally, pain assessment in cats benefits from established tools such as the Feline Grimace Scale, which provides a validated framework for evaluating facial expressions of pain in cats^58^. Overall, the results underscore the sensitivity and impact of landmark-based emotion classification to preprocessing choices such as alignment and augmentation and the need for optimised pipelines when working with small or noisy datasets^24,34^.

The predictive validity of the manual behaviour coding clearly outperformed the landmark-based approach, similar to the results of Austin et al.^42^. This substantial performance gap may be attributed in part to the limited size and variability of the landmark dataset, which likely reduced the model’s ability to generalise across subjects^34,37,42^. More fundamentally, manual behaviour coding captures semantically rich, observer-interpretable features that reflect both facial and full-body signals, but can be subject to human interpretation^1,13,24^. These include contextual behaviours such as posture, movement, and orientation, information not accessible through facial landmarks alone. Landmark-only models, though promising for automation, offer a scalable, low-cost solution but are inherently constrained by their narrow focus and current limitations in resolution and expressiveness, particularly when trained on small or limited datasets^30,33^. Possibly, combining automated facial landmark analysis with automated tail and posture analysis could increase the models’ validity in the future.

These findings reflect a tactical trade-off between the accuracy and interpretability of manual coding and the scalability and efficiency of automated methods. While manual annotation remains the gold standard for detailed emotional assessment, it is time-consuming and resource-intensive. In contrast, machine learning models—particularly those based on facial landmarks—offer the potential for rapid, scalable analysis but currently lack the nuanced understanding provided by full-body, observer-coded behaviours. Striking a balance between these approaches, or integrating them, could be key to advancing practical tools for real-time animal welfare monitoring.

In conclusion, the study demonstrates that firework-related fear in dogs could be classified with near-perfect accuracy based on manually coded expressions—substantially outperforming earlier results by Bremhorst et al.^25^ for predicting positive and negative situations based on individual DogFACS action units. This highlights the strength of machine learning in revealing consistent and discriminative expression patterns associated with distinct emotional states in dogs. These findings underscore the potential of validated behavioural indicators in advancing automated welfare assessment. Future development of the landmark approach could further enhance accuracy and generalizability by integrating additional features such as body posture or tail position, offering a multi-modal emotion detection. This combination could be a promising step toward automated welfare monitoring in these situations.

### Limitations of the study

The study’s small sample size and variable video quality are notable limitations, particularly for the landmark-based models, which rely on clear, stable facial imagery. The data set was highly challenging, as owners filming their own dog in their home setting led to different video angles, in part, very poor resolution and quality; and dogs were often only partially in view, e.g. when hiding under furniture. Additionally, the presence of owners during recordings may have influenced the dogs’ behaviour. For example, the obtained results could be different if data from other sources were used as a training/test set (e.g., dogs experiencing fireworks without the presence of the owner or outdoors, rather than inside), which could impact generalizability across firework contexts.

## Resource availability

### Lead Contact

For further information, please contact Jiao-Ling Appels

### Materials availability

This study did not generate new unique materials.

### Data and code availability

The dataset generated and analysed during the current study is available at Mendeley Data (https://doi.org/10.17632/swk97wxzzg.1). The original video recordings can be obtained from the corresponding author of Gähwiler et al. (2020) upon reasonable request. All analysis scripts used in this study are included in the Mendeley Data repository. Any other information required to reanalyse the data reported in this paper is available from the lead contact upon request.

## Acknowledgements

We thank all the dog caretakers who provided videos of their dogs.

## Ethics Declarations

The dataset used in this study was previously collected under ethical approval from the Veterinary Office of the Canton of Bern, Switzerland (Licence number BE28/17). The current study protocol was reviewed by the Ethical Committee of the University of Haifa, which determined that no additional approval was necessary.

## Authors Contributions

Conceptualisation, J.A., S.R., G.M.; Resources, S.R.; Methodology, S.R., G.M., A.Z.; formal analysis, J.A., G.M.; Visualisation, J.A., S.R.; Validation, A.Z.; Supervision, S.R., G.M., A.Z.; Writing—original draft, J.A.; writing—review and editing, J.A., S.R., G.M., A.Z.; all authors have read and agreed to the published version of the manuscript

## STAR★Methods

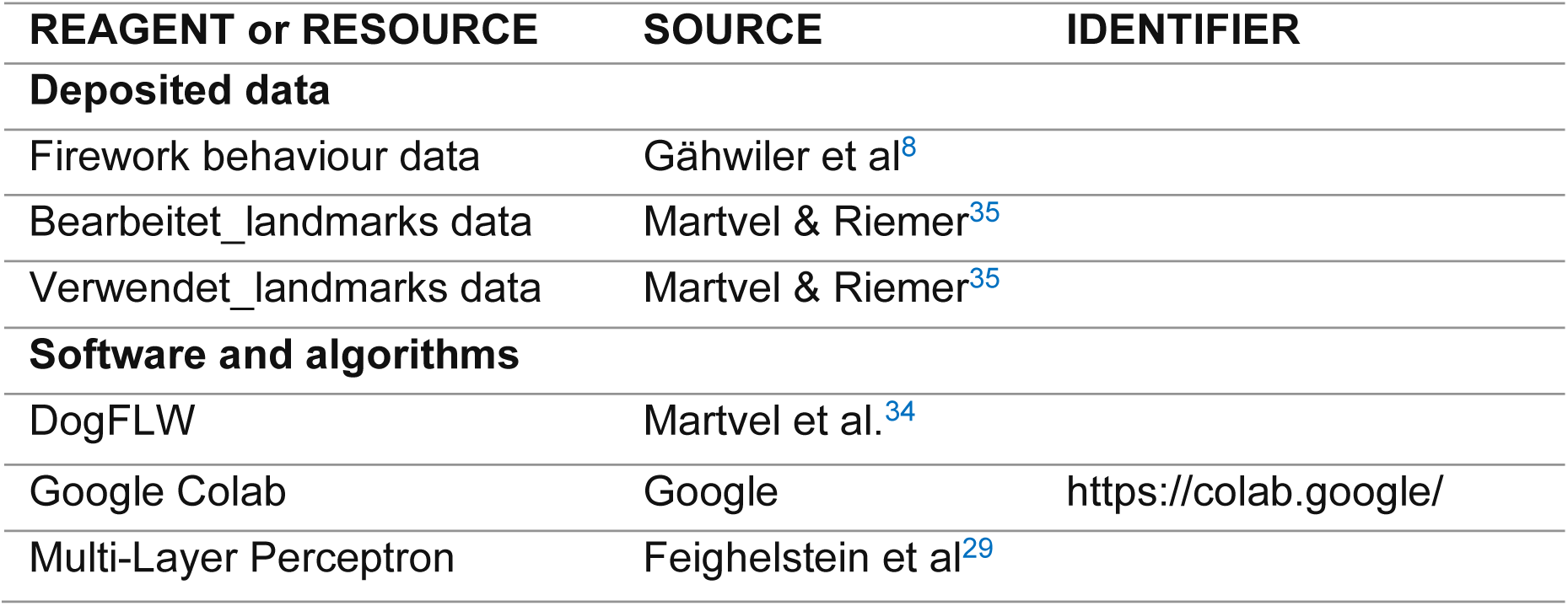
Key resources table.

## Data collection

### Videos

This study used a previously collected dataset of video recordings from 36 dogs (17 mixed breeds, 19 purebreds; 18 males, 18 females)^8^. Participating dog owners were recruited via social media, cynological associations, and dog sports groups. They took videos of their dog in a real-life firework situation (New Year’s Eve) and in a non-firework situation on a different evening. Owners were instructed to film one dog at a time and aim to capture the whole body on the video. For welfare reasons, owners were asked to behave normally—for example, by petting, feeding or calming the dog in their own way—during the firework situation. The videos were standardised for manual coding over the first three minutes. Using Solomon Coder, a blinded coder annotated behaviours based on a behavioural ethogram, including both whole-body (e.g., tail, posture, sitting) and facial behaviours (e.g., ear position, lip-licking, blinking).

Point sampling was used for scoring body, tail and ear postures and duration behaviours such as hiding or moving. Absolute durations were coded for time spent panting, sleeping or not visible (e.g., head, eyes or body out of view), and events such as lip licking and blinking were coded as frequencies (Table S4). Inter-rater reliability was established by an independent second coder on 46% of the videos (the first 60 seconds of 20 randomly selected videos comprising 10 fireworks and 10 control videos). Additionally, an extra reliability coding was done for rarer behaviours—panting and hiding—based on 13 non-randomly selected videos. The inter-rater reliability was good, with Cronbach’s α ≥ 0.70. In the original paper by Gähwiler et al.^8^, Wilcoxon signed-rank tests showed significant differences between the firework condition and the control condition in ear position score and the durations of locomotion and panting. The effects of vocalisations, blinking, and hiding became nonsignificant after sequential Bonferroni correction. In that study, ear position was assessed at the base of the ear and scored from 1-5 (1 = turned as far backwards as possible, 5 = directed forward, 3 = neutral position for the dog). Posture and tail position were likewise scored along a scale from 1-5, but were excluded from the analysis in the original paper because many dogs were lying during the control condition, so that posture and tail position were scored as NA. These previously reported results provide the statistical reference against which the current machine learning analysis was compared.

### Landmark detection and processing

The landmarks were automatically detected on all videos with an Ensemble Landmark Detector model^30^, trained on the DogFLW^34,35^.

### Evaluation metrics

All analyses were performed in Python (Google Colab) on a system equipped with an Intel Xeon CPU (2.2GHz), 13GB RAM, and an NVIDIA Tesla K80 GPU with 12GB memory. For both approaches (manual coding vs. landmarks), all models were trained and tested on the same dataset-behavioural counts in Approach 1 & landmark coordinate sets in Approach 2—using accuracy, precision, recall, and the F1-score^57,59^.

Accuracy measures the overall correctness of a model (the proportion of all images—both firework and non-firework—that were correctly classified); precision is the proportion of true positives (how many of the images predicted as ‘Firework’ truly belonged to firework situations); recall is correctly identified positives (how many of the actual firework images the model successfully identified); F1-score balances precision and recall (useful when the number of firework and non-firework images is imbalanced)^29^. ROC AUC is the area under the receiver operating characteristic curve^46^. This value shows the relation between true positives and false positives and therefore reflects the discriminative power and generalizability of a classification model^37,46^.

## Approach 1: Manually coded behavioural data

### Preprocessing

The dataset used for model training contained behavioural counts and durations coded from video observations (Document S1). These counts were derived from the original ethogram (Table S4). From this ethogram, a subset of 27 behaviours (Table S5) was chosen and used as predictors. Thus, the machine learning models were trained and tested with the coded dataset (Document S1), using the 27 selected behaviours (Table S5) as input features and the experimental conditions (firework vs. non-firework) as the groups for the classifier models. The dataset was split into 80% training and 20% testing using Scikit-learn’s *train_test_split* function^59^.

### Data processing

The data processing consisted of 5 steps (Figure 3). First, three machine learning models were trained for classification: Random Forest (RF) with 100 decision trees, a maximum depth of 10 to prevent overfitting, and Gini impurity for node splitting, with up to 35 features per split to reduce tree correlation. XGBoost (XGB) applied gain-based evaluation for dynamic feature importance. The Decision Tree (DT) model served as a baseline, providing a simpler, interpretable classification. Libraries used included Pandas, NumPy, Scikit-learn, XGBoost, Matplotlib, and SHAP. All hyperparameters were chosen empirically through a grid search.

**Figure 3:**
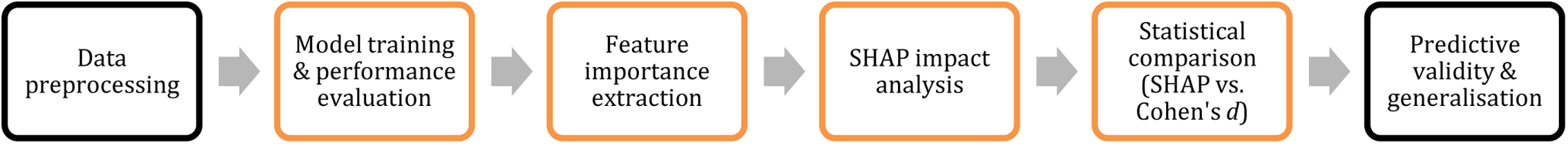
Process of the manual behavioural coding approach.

To determine the key behaviours driving model predictions, feature importance scores were extracted from all three models, following the top 5 best features of all models^60^. SHAP (SHapley Additive exPlanations) was used to visualise each feature’s contribution in the most accurate model^61,62^, providing interpretability in classifying firework-induced fear responses.

Additionally, the predictive power of machine learning models was compared to traditional statistical methods. Cohen’s *d* was used to assess effect size in the manual coding results^8^, and findings were evaluated alongside SHAP impact scores to determine unison between statistical significance and model-driven feature importance.

Moreover, to evaluate the predictive validity and generalizability of behavioural indicators of firework fear, several cross-validation strategies were implemented. First, an exploratory phase was conducted to compare the predictive performance of different feature sets. This included five configurations of features using RF with stratified k-fold cross-validation (k = 5 and k = 10): (1) top features based on raw feature importance, (2) top SHAP-ranked features, (3) SHAP-selected features with two additional top-ranking indicators, (4) a modified version of (3) including a different combination of extra features, and (5) the top 10 features by importance. A standard 10-fold cross-validation (CV) approach was used to benchmark model performance using both SHAP-selected features and the full set of behavioural indicators. The best-performing configuration, SHAP-selected features with tenfold cross-validation, was selected for further analysis.

Finally, to assess whether the predictive models generalize to unseen individuals, two subject-aware strategies were applied: (1) Leave-One-Subject-Out Cross-Validation (LOSO-CV), where each fold held out both conditions (firework and control) for a single dog, and (2) K-fold subject-based CV, where dogs were grouped into four folds (k ≈ number of dogs / 10) with no subject overlap between training and test sets. The dogs selected for the test set were included in all models and chosen to maximise variation and generalizability. Selection criteria included diversity in ear type (floppy vs. pointy), coat length (short vs. long), coat colour (light vs. dark), and body size (small vs. large). All models were evaluated using accuracy, precision, recall, F1 score, and ROC AUC. These validation methods were applied to models trained using both the full behavioural dataset and a reduced feature set comprising the most predictive indicators identified through SHAP. By testing multiple configurations, the study aimed to distinguish between raw model accuracy and true generalisation across individual animals.

## Approach 2: Landmark–based analysis

### Preprocessing

While approach 1 incorporated whole-body expressions, approach 2 isolated expression via facial landmarks (contouring of the ears, snout and mouth). The input for this approach consisted of data frames with facial landmark coordinate data extracted from videos of individual dogs in both conditions: firework and non-firework (control) (Document S2). Each data frame contained landmark data from every odd-numbered frame, as consecutive frames often showed minimal variation. There were 46 landmarks on the face of each dog: five for each ear; four for each eye; and the rest for the nose/mouth region and the base of the ears^34^ (Figure 4). The percentage of frames without landmarks was calculated for each video to assess the data quality in terms of informative frames. Videos with fewer than 50% of frames in which the face was successfully detected were excluded. From each remaining video, a fixed number of “good” frames (3, 5 or 10) were automatically and randomly selected using a Python script, resulting in a reproducible filtered landmark dataset. For example, the 3-frame-per-video dataset consisted of 96 landmark frames across 32 videos from 25 dogs, with 51 firework frames and 45 non-firework frames.

**Figure 4:**
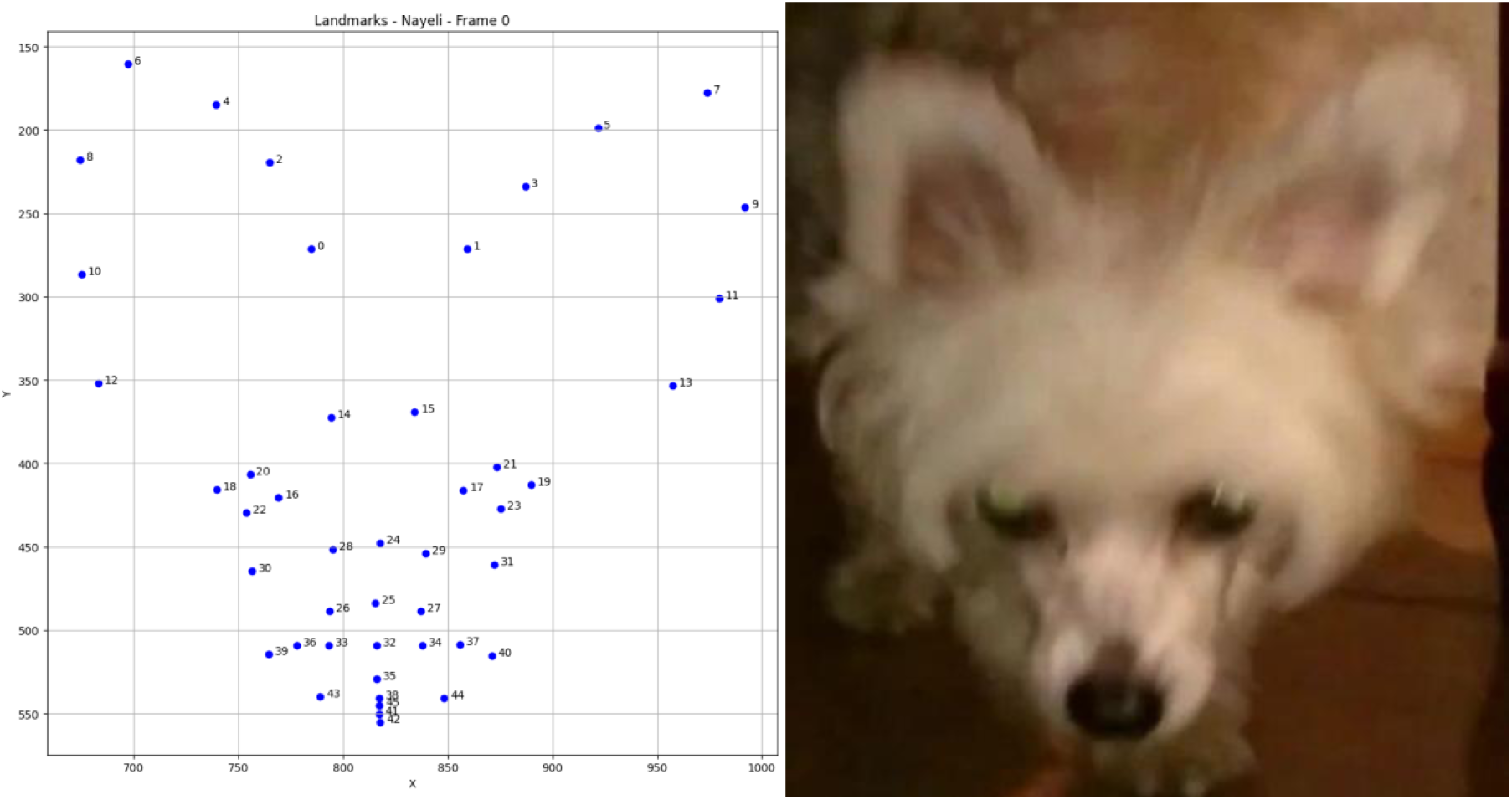
Facial landmark frame and the corresponding video screenshot frame.

### Ablation study design

To investigate the influence of preprocessing and dataset composition, an ablation study was conducted. Models were trained under systematically varied conditions. All configurations were run on both the full set of 32 videos from 25 dogs and on a balanced subset of 7 dogs (with equal numbers of firework and control videos). The analysis was run for each subject set with different frame sampling (3, 5 or 10 frames per video).

### Processing pipeline

Following Feighelstein et al.^29^, facial landmarks were processed in two main steps: alignment and data augmentation (Figure 5). First, facial alignment was applied to centre the landmark coordinates using the outer eye landmark points as reference anchors, defined from the dog’s perspective (landmark 18 = anatomical right eye, landmark 19 = left eye). Second, data augmentation—from existing data, generating new data and adding that to the training set—was performed via Gaussian noise injection to improve generalisation and model robustness. This is achieved by applying small, random perturbations to the original landmark coordinates. Specifically, each coordinate (X, Y) from the annotated landmarks was multiplied by a random value drawn from a normal distribution with a mean of 1 and a standard deviation of 0.0005. This process introduced subtle variations in the input data, simulating natural variability in automated detection, without significant morphological changes. The full pipeline transforms each annotated image—containing 46 facial landmarks—into a feature vector of all x and y coordinates^35^.

**Figure 5:**
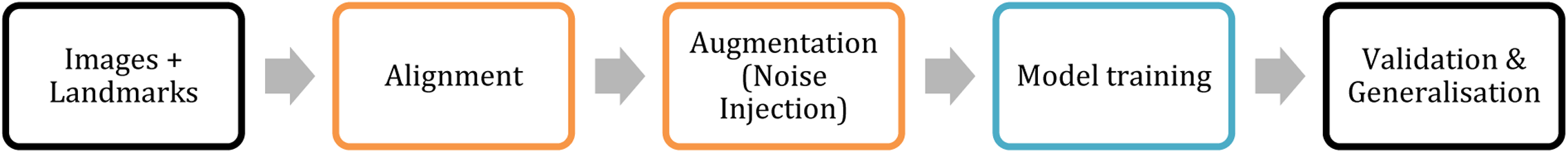
Process of the landmark-based approach.

### Model training

We implemented a Multi-Layer Perceptron (MLP) neural network comprising an input layer with 92 neurons, corresponding to the x and y coordinates of the 46 facial landmarks. The network architecture includes three hidden layers, which were based on the validated approach of Feighelstein et al.^29^: the first and second layers each contain 100 neurons with ReLU activation functions, while the third layer contains 500 neurons, also using ReLU activation. The output layer consists of two neurons with a Softmax activation function to support binary classification. The model was trained for 10 epochs using the Adam optimiser with a learning rate of 0.01 and a batch size of 32. During each epoch, the training data were normalised using standard scaling and augmented as described previously. Given the small dataset size (max. 320 frames), extensive hyperparameter tuning was avoided to reduce the risk of overfitting. Instead, the final model was selected based on the lowest validation loss observed during training, ensuring optimal generalisation performance.

To assess the impact of preprocessing techniques, the same MLP architecture was trained under four different conditions, varying in the use of facial alignment and data augmentation. These conditions included: (1) no alignment or augmentation, (2) alignment only, (3) augmentation only, and (4) both alignment and augmentation. This allowed evaluation of how these preprocessing steps influenced classification performance.

This approach was chosen instead of Random Forest or XGBoost due to the spatial nature of the landmark data. Tree-based models are well-suited for behavioural frequency data, as used in Approach 1. In contrast, the coordinate data from this approach required a model capable of capturing spatial patterns, for which MLPs are more appropriate. MLPs can capture nonlinear interactions between all landmark positions, allowing them to learn spatial patterns that are essential for distinguishing facial expressions^63^.

### Validation

As a validation method, we used leave-one-subject-out (LOSO) cross-validation with no subject overlap. Due to the relatively low numbers of dog videos (n=32 or 14) and frames in the dataset, following the stricter method is more appropriate. In our case, this means that we repeatedly left-out one subject from training and validation. We trained on 80% of the remaining subjects, validated with the other 20% of subjects and tested on the one left out subject. By separating the subjects used for training, validation and testing, respectively, we enforced generalisation to unseen subjects and ensured that no specific features of an individual are used for classification. To enable consistent comparison across models and dataset configurations, we defined a fixed test set consisting of three dogs. This number was chosen to accommodate the smallest dataset (n = 7), where only four subjects remained for training and validation. The selected test subjects varied in ear shape (floppy vs. pointy), coat colour and length (light/dark, short/long), and body size to ensure morphological diversity and visual generalizability.

## Supplemental information (1)

**Document S1 and S2. Tables S1, S2, S3, S4 and S5.**

## Supplemental documents

Document S1: Manually coded behaviour dataset

Firework_Video_Data_cleaned.xlsx

Document S2: Two used landmark data sets bearbeitet_landmarks & verwendet_landmarks

### Supplemental tables

**Table S1:** Model metrics and feature importance.

| Random Forest model |  |  |  |  |  |  |
| --- | --- | --- | --- | --- | --- | --- |
| Model metrics |  |  |  |  | Feature importance top 5 |  |
|  | Precision | Recall | F1-score | Support | Behaviour | Importance (%) |
| **Firework** | 0.89 | 0.80 | 0.84 | 10 | Ear 1 (low) | 19,875166 |
| **Control** | 0.67 | 0.80 | 0.73 | 5 | Ear 2 | 11,583891 |
|  |  |  |  |  | Blink/min | 10,066911 |
| **Accuracy** |  |  | 0.80 | 15 | Ear 3 | 7,351706 |
| **Macro avg** | 0.78 | 0.80 | 0.78 | 15 | Move (%) | 6,832791 |
| **Weighted avg** | 0.81 | 0.80 | 0.80 | 15 |  |  |
| XGBoost model |  |  |  |  |  |  |
| Model metrics |  |  |  |  | Feature importance top 5 |  |
|  | Precision | Recall | F1-score | Support | Behaviour | Importance (%) |
| **Firework** | 0.75 | 0.60 | 0.67 | 10 | Posture 2 | 43,917801 |
| **Control** | 0.43 | 0.60 | 0.50 | 5 | Ear 1 (low) | 22,944616 |
|  |  |  |  |  | Move (%) | 10,071712 |
| **Accuracy** |  |  | 0.60 | 15 | Ear 3 | 3,564093 |
| **Macro avg** | 0.59 | 0.60 | 0.58 | 15 | Posture 3 | 3,485780 |
| **Weighted avg** | 0.64 | 0.60 | 0.61 | 15 |  |  |
| Decision Tree model |  |  |  |  |  |  |
| Model metrics |  |  |  |  | Feature importance top 5 |  |
|  | Precision | Recall | F1-score | Support | Behaviour | Importance (%) |
| **Firework** | 0.78 | 0.70 | 0.74 | 10 | Ear 1 (low) | 31,121187 |
| **Control** | 0.50 | 0.60 | 0.55 | 5 | Posture 2 | 26,239053 |
|  |  |  |  |  | Lying (%) | 9,547146 |
| **Accuracy** |  |  | 0.67 | 15 | Ear 3 | 8,615800 |
| **Macro avg** | 0.64 | 0.65 | 0.64 | 15 | Yawn/min | 6,629963 |
| **Weighted avg** | 0.69 | 0.67 | 0.67 | 15 |  |  |

**Table S2:** Feature Importance (current study) vs. Effect Size (Cohen’s *d*) from Gähwiler et al. (2020)

| Machine Learning: random forest |  |  |  | Statistics: Cohen's *d* |  |
| --- | --- | --- | --- | --- | --- |
| Top 6 best features |  |  |  | Effect size |  |
| **1** | Ear 1 (low) | 19,875166 | % | Ear position* | 0.68 |
| **2** | Ear 2 | 11,583891 | % | Locomotion* | 0.54 |
| **3** | Blink/min | 10,066911 | % | Panting* | 0.45 |
| **4** | Ear 3 | 7,351706 | % | Vocalisation | 0.40 |
| **5** | Move (%) | 6,832791 | % | Blinking | 0.37 |
| **6** | Posture 2 | 5,482166 | % | Hiding | 0.37 |

\*significant variables

**Table S3:**
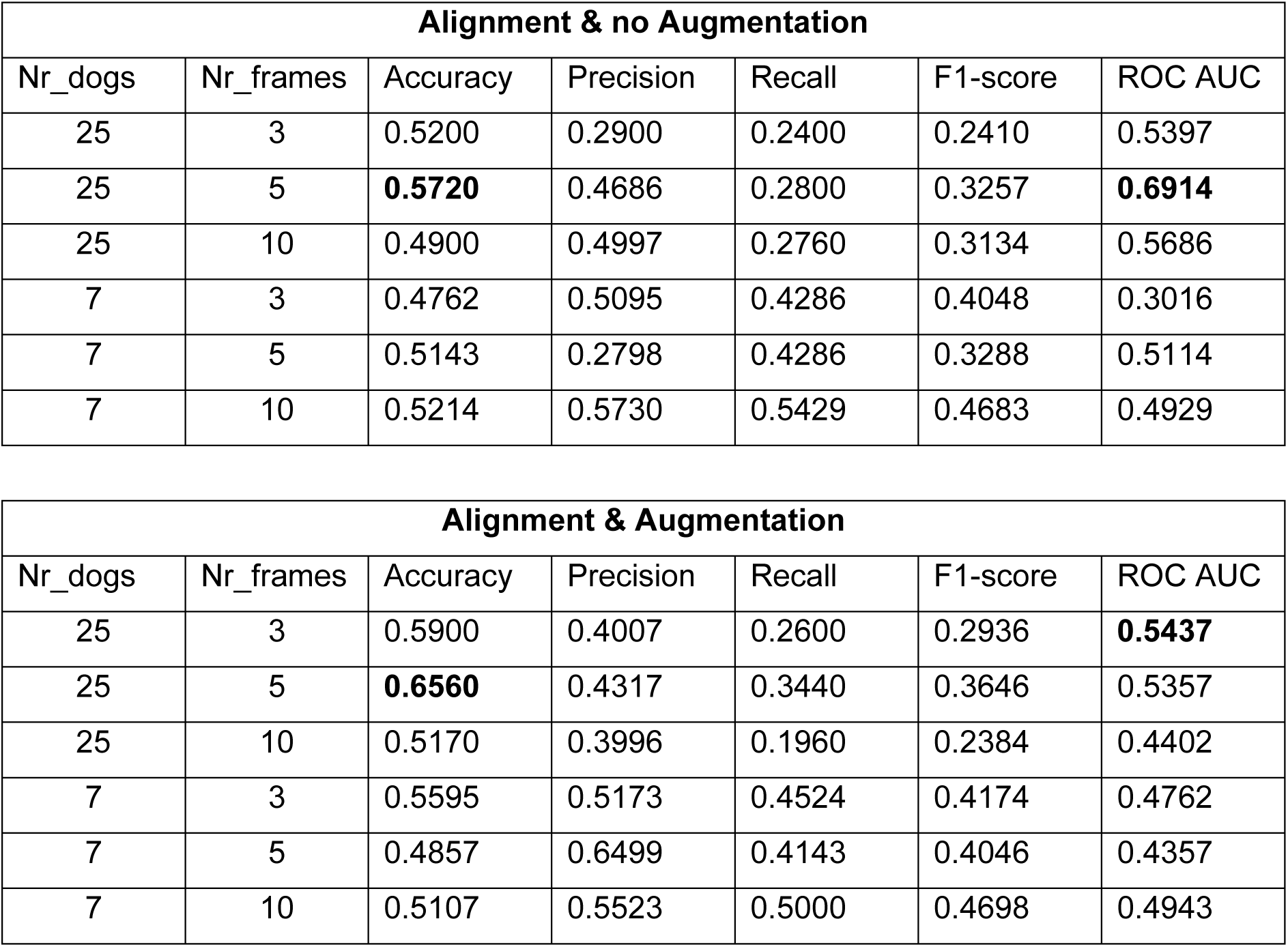
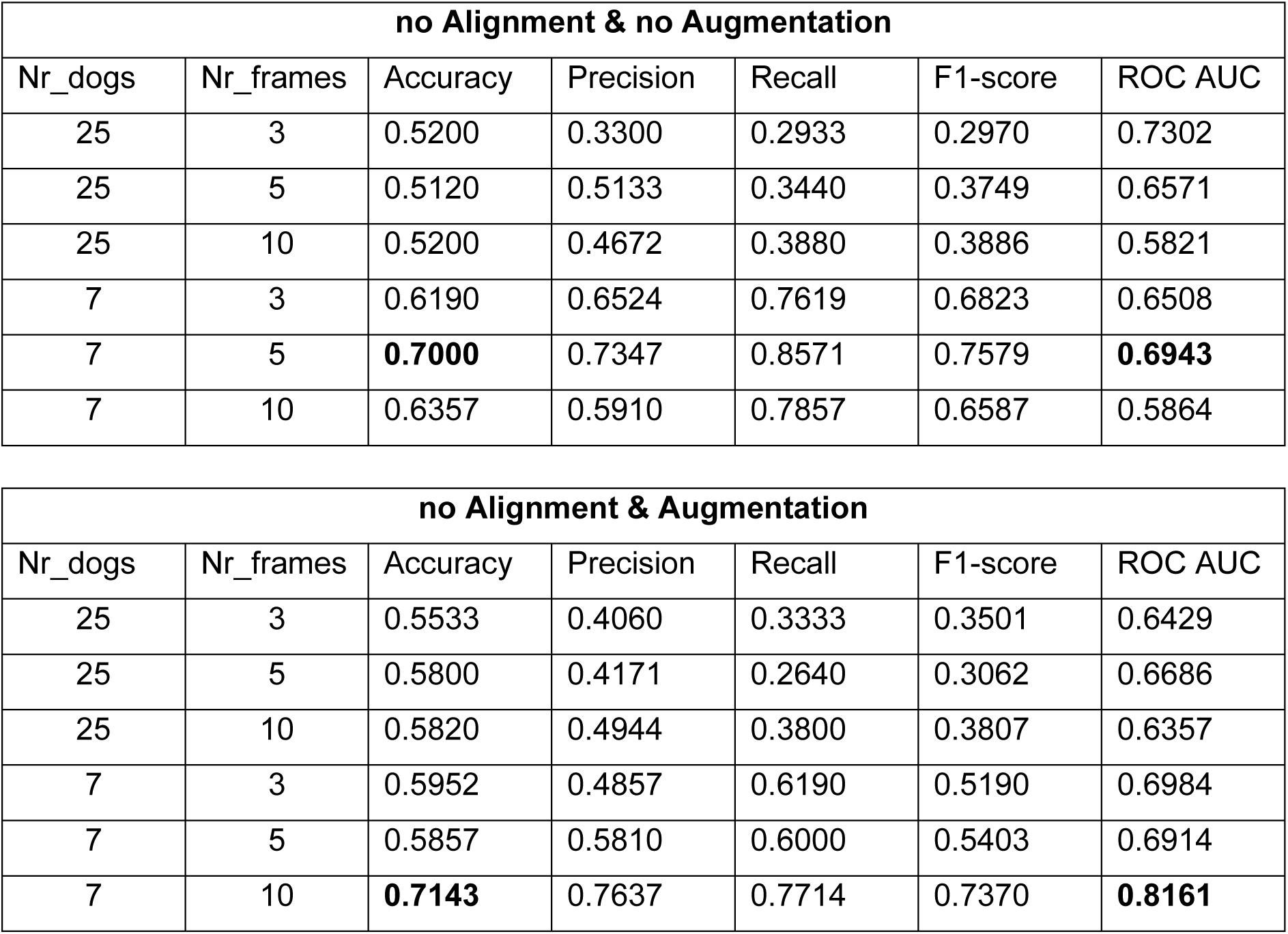
Evaluation metrics of all landmark approaches and configurations. Best values in bold.

| Alignment & no Augmentation |  |  |  |  |  |  |
| --- | --- | --- | --- | --- | --- | --- |
| Nr_dogs | Nr_frames | Accuracy | Precision | Recall | F1-score | ROC AUC |
| 25 | 3 | 0.5200 | 0.2900 | 0.2400 | 0.2410 | 0.5397 |
| 25 | 5 | **0.5720** | 0.4686 | 0.2800 | 0.3257 | **0.6914** |
| 25 | 10 | 0.4900 | 0.4997 | 0.2760 | 0.3134 | 0.5686 |
| 7 | 3 | 0.4762 | 0.5095 | 0.4286 | 0.4048 | 0.3016 |
| 7 | 5 | 0.5143 | 0.2798 | 0.4286 | 0.3288 | 0.5114 |
| 7 | 10 | 0.5214 | 0.5730 | 0.5429 | 0.4683 | 0.4929 |

| Alignment & Augmentation |  |  |  |  |  |  |
| --- | --- | --- | --- | --- | --- | --- |
| Nr_dogs | Nr_frames | Accuracy | Precision | Recall | F1-score | ROC AUC |
| 25 | 3 | 0.5900 | 0.4007 | 0.2600 | 0.2936 | **0.5437** |
| 25 | 5 | **0.6560** | 0.4317 | 0.3440 | 0.3646 | 0.5357 |
| 25 | 10 | 0.5170 | 0.3996 | 0.1960 | 0.2384 | 0.4402 |
| 7 | 3 | 0.5595 | 0.5173 | 0.4524 | 0.4174 | 0.4762 |
| 7 | 5 | 0.4857 | 0.6499 | 0.4143 | 0.4046 | 0.4357 |
| 7 | 10 | 0.5107 | 0.5523 | 0.5000 | 0.4698 | 0.4943 |

| no Alignment & no Augmentation |  |  |  |  |  |  |
| --- | --- | --- | --- | --- | --- | --- |
| Nr_dogs | Nr_frames | Accuracy | Precision | Recall | F1-score | ROC AUC |
| 25 | 3 | 0.5200 | 0.3300 | 0.2933 | 0.2970 | 0.7302 |
| 25 | 5 | 0.5120 | 0.5133 | 0.3440 | 0.3749 | 0.6571 |
| 25 | 10 | 0.5200 | 0.4672 | 0.3880 | 0.3886 | 0.5821 |
| 7 | 3 | 0.6190 | 0.6524 | 0.7619 | 0.6823 | 0.6508 |
| 7 | 5 | **0.7000** | 0.7347 | 0.8571 | 0.7579 | **0.6943** |
| 7 | 10 | 0.6357 | 0.5910 | 0.7857 | 0.6587 | 0.5864 |

| no Alignment & Augmentation |  |  |  |  |  |  |
| --- | --- | --- | --- | --- | --- | --- |
| Nr_dogs | Nr_frames | Accuracy | Precision | Recall | F1-score | ROC AUC |
| 25 | 3 | 0.5533 | 0.4060 | 0.3333 | 0.3501 | 0.6429 |
| 25 | 5 | 0.5800 | 0.4171 | 0.2640 | 0.3062 | 0.6686 |
| 25 | 10 | 0.5820 | 0.4944 | 0.3800 | 0.3807 | 0.6357 |
| 7 | 3 | 0.5952 | 0.4857 | 0.6190 | 0.5190 | 0.6984 |
| 7 | 5 | 0.5857 | 0.5810 | 0.6000 | 0.5403 | 0.6914 |
| 7 | 10 | **0.7143** | 0.7637 | 0.7714 | 0.7370 | **0.8161** |

**Table S4:**
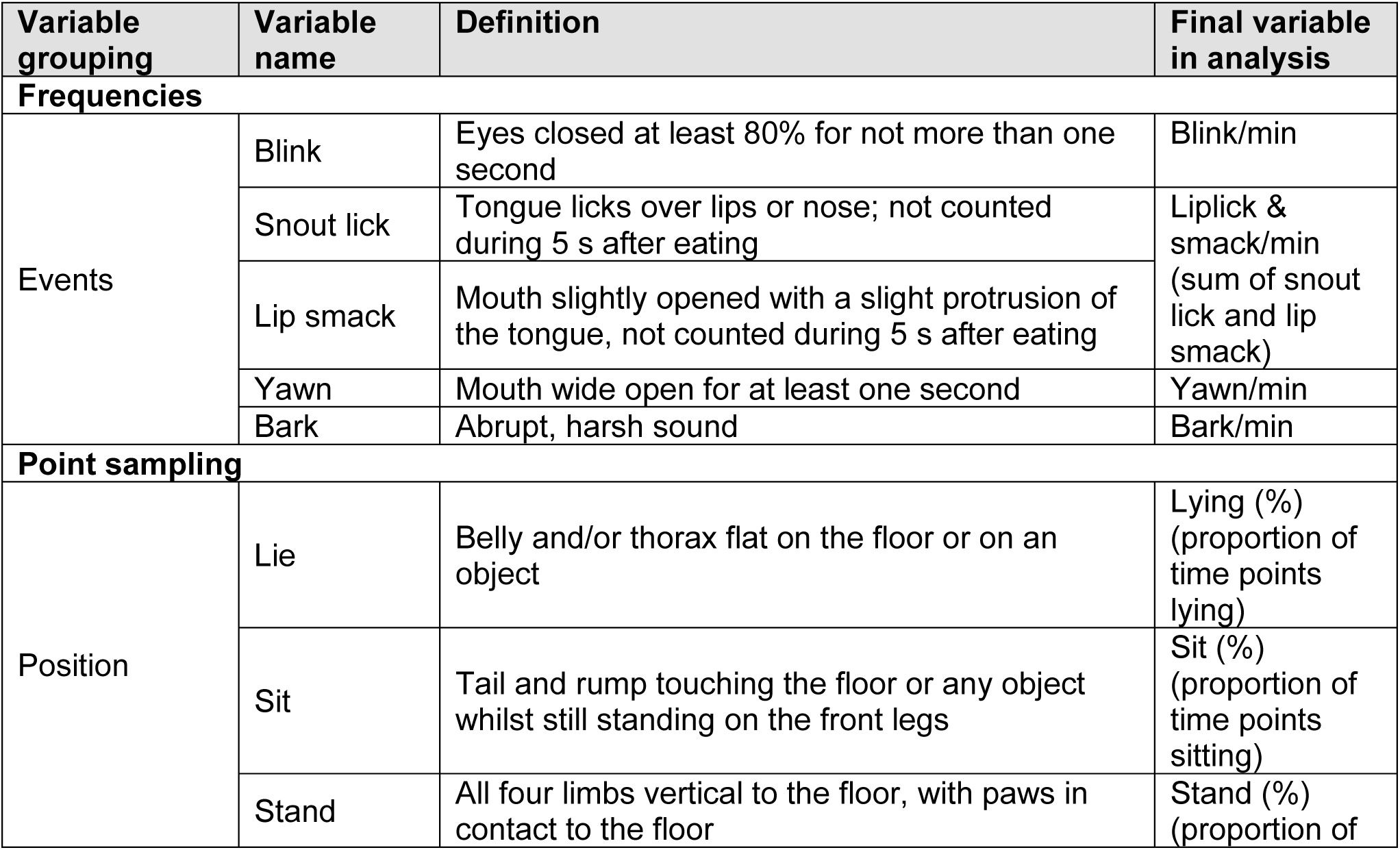
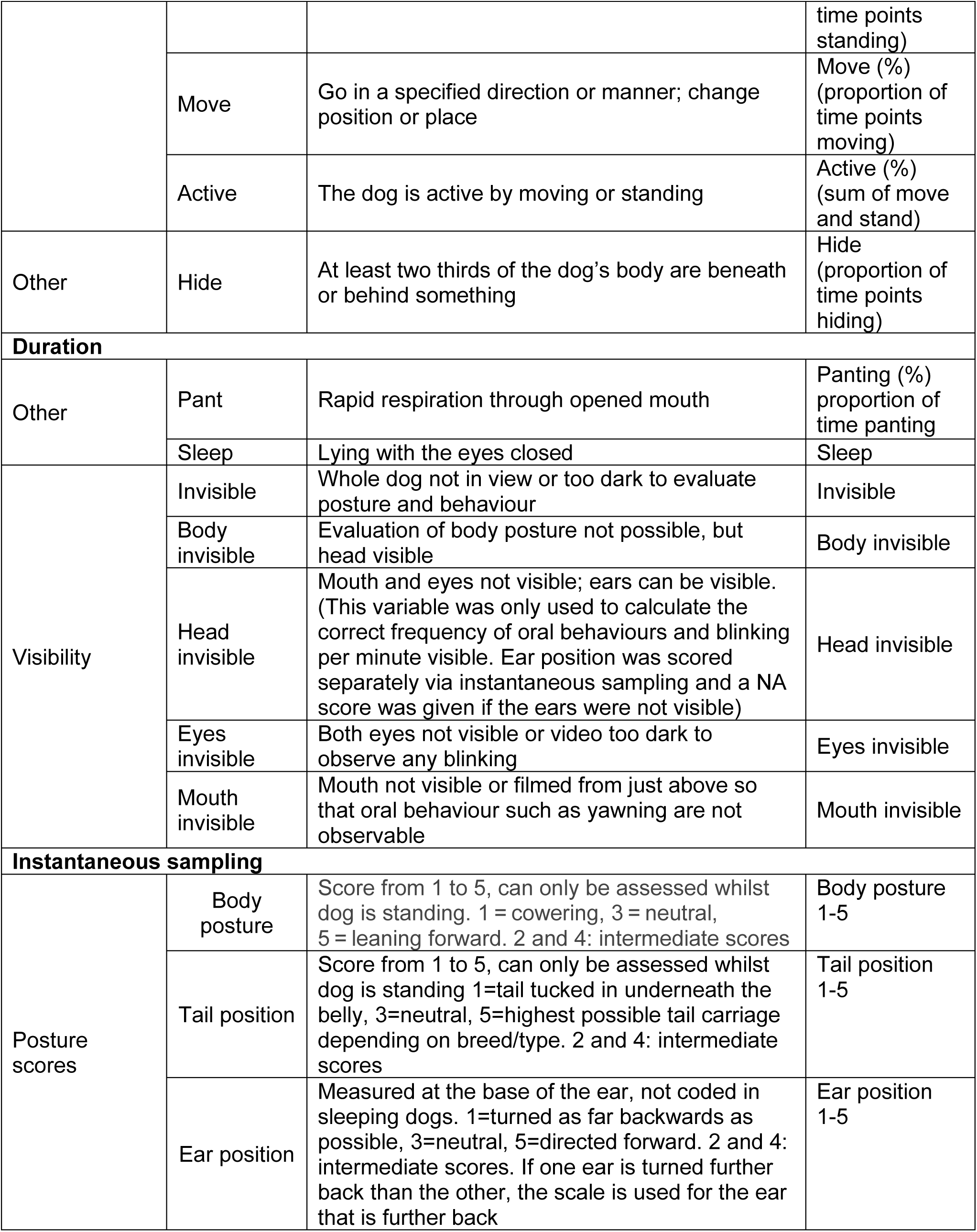
Full ethogram used for behaviour coding.

| Variable grouping | Variable name | Definition | Final variable in analysis |
| --- | --- | --- | --- |
| **Frequencies** |  |  |  |
| Events | Blink | Eyes closed at least 80% for not more than one second | Blink/min |
|  | Snout lick | Tongue licks over lips or nose; not counted during 5 s after eating | Liplick & smack/min<br>(sum of snout lick and lip smack) |
|  | Lip smack | Mouth slightly opened with a slight protrusion of the tongue, not counted during 5 s after eating |  |
|  | Yawn | Mouth wide open for at least one second | Yawn/min |
|  | Bark | Abrupt, harsh sound | Bark/min |
| **Point sampling** |  |  |  |
| Position | Lie | Belly and/or thorax flat on the floor or on an object | Lying (%)<br>(proportion of time points lying) |
|  | Sit | Tail and rump touching the floor or any object whilst still standing on the front legs | Sit (%)<br>(proportion of time points sitting) |
|  | Stand | All four limbs vertical to the floor, with paws in contact to the floor | Stand (%)<br>(proportion of time points standing) |
|  | Move | Go in a specified direction or manner; change position or place | Move (%) (proportion of time points moving) |
|  | Active | The dog is active by moving or standing | Active (%) (sum of move and stand) |
| Other | Hide | At least two thirds of the dog's body are beneath or behind something | Hide (proportion of time points hiding) |
| **Duration** |  |  |  |
| Other | Pant | Rapid respiration through opened mouth | Panting (%) proportion of time panting |
|  | Sleep | Lying with the eyes closed | Sleep |
| Visibility | Invisible | Whole dog not in view or too dark to evaluate posture and behaviour | Invisible |
|  | Body invisible | Evaluation of body posture not possible, but head visible | Body invisible |
|  | Head invisible | Mouth and eyes not visible; ears can be visible. (This variable was only used to calculate the correct frequency of oral behaviours and blinking per minute visible. Ear position was scored separately via instantaneous sampling and a NA score was given if the ears were not visible) | Head invisible |
|  | Eyes invisible | Both eyes not visible or video too dark to observe any blinking | Eyes invisible |
|  | Mouth invisible | Mouth not visible or filmed from just above so that oral behaviour such as yawning are not observable | Mouth invisible |
| **Instantaneous sampling** |  |  |  |
| Posture scores | Body posture | Score from 1 to 5, can only be assessed whilst dog is standing. 1 = cowering, 3 = neutral, 5 = leaning forward. 2 and 4: intermediate scores | Body posture 1-5 |
|  | Tail position | Score from 1 to 5, can only be assessed whilst dog is standing 1=tail tucked in underneath the belly, 3=neutral, 5=highest possible tail carriage depending on breed/type. 2 and 4: intermediate scores | Tail position 1-5 |
|  | Ear position | Measured at the base of the ear, not coded in sleeping dogs. 1=turned as far backwards as possible, 3=neutral, 5=directed forward. 2 and 4: intermediate scores. If one ear is turned further back than the other, the scale is used for the ear that is further back | Ear position 1-5 |

**Table S5:** Behaviour subset used in ML models.

| Nr. | Variable name | Variable in dataset | Definition |
| --- | --- | --- | --- |
| 1 | Ear 1 (low) | Ear_1_low | Measured at the base of the ear, not coded in sleeping dogs. 1 = turned as far backwards as possible, 3 = neutral, 5 = directed forward. 2 and 4: intermediate scores. |
| 2 | Ear 2 | Ear_2 | ↓ |
| 3 | Ear 3 | Ear_3 |  |
| 4 | Ear 4 | Ear_4 |  |
| 5 | Ear 5 | Ear_5 |  |
| 6 | Tail 1 | Tail_1 | Score from 1 to 5, can only be assessed whilst dog is standing. 1 = tail tucked in underneath the belly, 3 = neutral, 5 = highest possible tail carriage depending on breed/type. 2 and 4: intermediate scores |
| 7 | Tail 2 | Tail_2 | ↓ |
| 8 | Tail 3 | Tail_3 |  |
| 9 | Tail 4 | Tail_4 |  |
| 10 | Tail 5 | Tail_5 |  |
| 11 | Posture 1 | Post_1 | Score from 1 to 5, can only be assessed whilst dog is standing. 1 = cowering, 3 = neutral, 5 = leaning forward. 2 and 4: intermediate scores |
| 12 | Posture 2 | Post_2 | ↓ |
| 13 | Posture 3 | Post_3 |  |
| 14 | Posture 4 | Post_4 |  |
| 15 | Posture 5 | Post_5 |  |
| 16 | Bark/min | Barking_min | Frequency of barking per minute |
| 17 | Hide | Hide | At least two thirds of the dog's body are beneath or behind something |
| 18 | Panting (%) | real_pant_percent | Percentage of panting from the total observations (panting & no panting) |
| 19 | Sleep | Sleep | Relaxed lying, eyes closed |
| 20 | Liplick & lipsmack/min | liplick_AND_smack_min | Sum of liplicks and smacks divided by total time visible |
| 21 | Blink/min | blink_min | Frequency of blinking per minute |
| 22 | Yawn/min | yawn_min | Frequency of yawning per minute |
| 23 | Move (%) | movepercent | Percentage of moving, calculated from the total number of position observations (move, stand, sit, lie, sleep) |
| 24 | Stand (%) | standpercent | Percentage of standing, calculated from the total number of position observations (move, stand, sit, lie, sleep) |
| 25 | Active (%) | activepercent | Sum of Move (%) and Stand (%) |
| 26 | Lying (%) | liepercent | Percentage of lying down, calculated from the total number of position observations (move, stand, sit, lie, sleep) |
| 27 | Sitting (%) | sitpercent | Percentage of sitting, calculated from the total number of position observations (move, stand, sit, lie, sleep) |

